# Development and Evaluation of a PCR Test System for the Detection of *Salmonella* spp. in Clinical and Epidemiological Materials

**DOI:** 10.64898/2026.06.24.734224

**Authors:** Duman Yessimseit, Altynai Kassenova, Beck Abdeliyev, Altyn Rysbekova, Zauresh Zhumadilova, Ziyat Abdel, Raikhan Mussagaliyeva, Tatyana Meka-Mechenko, Elmira Begimbayeva, Zhanar Nusipzhanova, Ayaulym Maksatova, Sanzhar Agzam, Gulmira Abdrassilova, Bakdaulet Kulbek, Oleg Reva, Aigul Abdirassilova

**Affiliations:** M. Aikimbayev National Scientific Center for Especially Dangerous Infections, 14 Zhakhanger St., Almaty A35P0K3, Kazakhstan; Al-Farabi Kazakh National University, 71 al-Farabi Ave., Almaty A15E3B4, Kazakhstan; Centre for Bioinformatics and Computational Biology, Department of Biochemistry, Genetics and Microbiology, University of Pretoria, Pretoria 0002, South Africa

**Keywords:** salmonellosis, *Salmonella*, PCR, real-time PCR, molecular diagnostics, *hilA*, TaqMan Assay, pathogen surveillance, food-borne infection, epidemiology

## Abstract

**Background:** Reliable detection of *Salmonella* remains a major challenge for public health surveillance and food safety due to the growing diversity of circulating serovars and the limitations of existing molecular targets. This study aimed to identify an optimal molecular target and develop a TaqMan real-time PCR assay for the detection of *Salmonella* spp.

**Methods:** Based on the results screening for *Salmonella* genes suitability as molecular markers, a TaqMan real-time PCR assay targeting the *hilA* gene was developed and validated. Analytical sensitivity, analytical specificity, and performance on bacterial isolates and artificially contaminated food samples were assessed.

**Results:** Among all candidate targets, *hilA* demonstrated the broadest coverage and was detected in all tested *Salmonella* isolates, including representatives of rare serological groups, whereas *invA* conventionally used for this pathogen detection, was absent in a subset of strains. The assay exhibited a limit of detection of 100 bacterial cells/mL and 100 fg/μL of genomic DNA. No cross-reactivity was observed with DNA from *Shigella flexneri*, *Shigella sonnei*, *Yersinia pestis*, *Y. pseudotuberculosis*, *Y. enterocolitica*, *Y. kristensenii*, *Bacillus anthracis*, *Vibrio cholerae*, or *Francisella tularensis*. The assay successfully detected *Salmonella* DNA in all artificially contaminated food samples tested. Evaluation using a collection of 25 bacterial isolates demonstrated positive amplification in all 24 confirmed *Salmonella* strains, while a strain initially identified by conventional bacteriology as *Salmonella* but subsequently confirmed by whole-genome sequencing as *Proteus mirabilis* yielded a negative result.

**Conclusions:** The *hilA* gene represents a highly conserved and reliable molecular target for the detection of *Salmonella* spp. The developed TaqMan real-time PCR assay demonstrated high analytical sensitivity, excellent specificity, and broad serovar coverage, supporting its application in laboratory detection of *Salmonella*, food safety monitoring, and epidemiological surveillance.

## 1. Introduction

Salmonellosis remains one of the most pressing problems in infectious disease pathology and public health worldwide. Among acute intestinal infections, diseases caused by bacteria of the genus *Salmonella* rank among the leading causes in terms of prevalence and medical-social significance [1-**Error! Reference source not found.**]. According to the World Health Organization, millions of cases of non-typhoidal salmonellosis are reported annually, resulting in substantial morbidity and mortality [4,5].

The causative agents of salmonellosis are characterized by a high pathogenic potential and considerable serological diversity. According to the Kauffmann-White classification, more than 2,500 serovars of *Salmonella* have been described, among which *S. enterica* Enteritidis and Typhimurium are of the greatest epidemiological importance [6]. In recent years, an increase has been observed in the proportion of rare *Salmonella* serotypes involved in both sporadic cases and outbreaks of foodborne infections [**Error! Reference source not found.**]. An additional concern is the spread of antibiotic-resistant and multidrug-resistant strains associated with both community-acquired and nosocomial infections [8].

According to official statistics, the incidence of salmonellosis has remained consistently high in recent years, while *S. enterica* Enteritidis and Typhimurium continue to dominate among circulating serotypes [9–11]. At the same time, the proportion of acute intestinal infections of unspecified etiology, particularly among children, is increasing, highlighting the need for improved laboratory diagnostic methods [12, **Error! Reference source not found.**].

The conventional bacteriological method, considered the gold standard for the diagnosis of salmonellosis, has several limitations, including the lengthy turnaround time, the requirement for isolation of a pure culture, biochemical identification procedures, and substantial laboratory infrastructure requirements [14]. Consequently, molecular genetic diagnostic methods, particularly real-time polymerase chain reaction (real-time PCR), have gained increasing importance due to their high sensitivity, specificity, and rapid turnaround time [14,15].

At present, a number of genetic targets have been proposed for the molecular diagnosis of *Salmonella* spp., including *invA*, *hilA*, *phoP*, *fimA*, *rpoS*, and *spvA*. However, published studies indicate that some molecular markers may be absent in certain *Salmonella* serovars, including representatives of rare serological groups, thereby limiting the universality of existing PCR-based assays [16,17]. In this context, the *hilA* gene, located within the SPI-1 pathogenicity island and responsible for regulating the expression of invasion-associated genes, is of particular interest. The high degree of conservation of the hilA gene among diverse *Salmonella* serovars makes it a promising molecular target for the universal PCR detection of salmonellosis pathogens [18,19].

Timely and accurate laboratory diagnosis of salmonellosis is of particular clinical importance in light of the rare but severe extra-intestinal complications of this infection reported in the literature [8], including osteomyelitis [20]. The development of these complications may be delayed in recognition or obscured in the absence of prompt etiological confirmation.

The aim of the present study was to identify an optimal molecular target and to develop and evaluate the performance of a real-time PCR assay for the detection of *Salmonella* spp. based on the detection of the marker gene *hilA* in clinical, food, and experimental samples.

## 2. Materials and Methods

### 2.1. Bacterial isolates and study materials

All procedures performed in this study complied with international and national ethical requirements for biomedical research and were approved by the Local Bioethics Committee of the M. Aikimbayev National Scientific Center for Especially Dangerous Infections (Protocol No. 2, dated 12 February 2024).

*Salmonella* strains used in this study were selected from the National Collection of Microorganisms (NCM) maintained at the M. Aikimbayev National Scientific Center for Especially Dangerous Infections (NSCEDI, https://nscedi.kz/en/) in Almaty, Kazakhstan. The isolates had been collected over different years and represented a variety of *Salmonella* serovars, including isolates belonging to rare serological groups. These strains were cultured on Endo and Ploskirev selective and differential diagnostic media. The cultures were incubated aerobically at 37°C for 18–24 h. Colony morphology was assessed according to standard bacteriological procedures [17].

Preliminary identification of the isolates was performed based on their cultural and biochemical characteristics. Serological identification was carried out using the slide agglutination test with a polyvalent adsorbed O-antiserum against *Salmonella* serogroups A–E, according to the manufacturer’s instructions. Serovar assignment was confirmed using specific O and H antisera in accordance with the Kauffmann-White-Le Minor scheme [21].

Serovar determination, whole-genome sequencing (WGS), and MLST typing of the selected strains were performed as part of a parallel study published separately [22]. Information on the source of isolation of the *S. enterica* strains, together with the results of serological and genetic typing, is presented in Table 1.

**Table 1.** Characteristics of the strains used in this study and the results of their serovar and genetic typing.

| Strain | Isolation source | Date of isolation | Region | Serovar name and type* | MLST |
| --- | --- | --- | --- | --- | --- |
| 2S | Animal | 02.07.1977 | Almaty region | *S. enterica* Paratyphi B 4:b:1,2 | ST86 |
| 5S | Not recorded | 15.03.1989 | Almaty region | *S. enterica* Infantis 7:r:1,5 | ST32 |
| 6S | Rabbit | 15.03.1989 | Almaty region | *S. enterica* Enteritidis 9:g,m:- | ST11 |
| 8S | Not recorded | 25.04.1963 | Kyrgyzstan | *S. enterica* Typhimurium 4:i:1,2 | ST328 |
| 10S | Animal | 25.11.2009 | Almaty region | *S. enterica* Dublin 9:g,p:- | ST10 |
| 13S | Not recorded | 25.11.2009 | Shymkent region | *S. enterica* Kottbus 8:e,h:1,5 | ST582 |
| 15S | Not recorded | 25.11.2009 | Shymkent city | *S. enterica* Typhi 9:d:- | ST1 |
| 16S | Human | 18.09.2025 | Almaty city | *S. enterica* Anatum | ST11 |
| 17S | Human | 07.12.2025 | Almaty city | *S. enterica* Enteritidis 9:g,m:- | ST11 |
| 18S<sup>†</sup> | Human | 26.10.2025 | Almaty city | *Proteus mirabilis* | ST490 |
| 20S | Human | 07.11.2025 | Almaty city | *S. enterica* Typhimurium 4:i:1,2 | ST376 |
| 23S | An aborted sheep fetus | 04.01.2024 | Karaganda region | *S. enterica* Choleraesuis 7:c:1,5 | ST68 |
| 24S | Horse (*Equus ferus caballus*) | 27.02.2012 | Almaty region | *S. enterica* Choleraesuis 7:c:1,5 | ST68 |
| 25S | Calf (*Bos taurus*) | 09.04.2013 | Almaty region | *S. enterica* Dublin 9:g,p:- | ST10 |
| 26S | Human | 28.06.2025 | Almaty city | *S. enterica* Typhimurium 4:i:1,2 | ST19 |
| 27S | Human | 19.09.2025 | Almaty city | *S. enterica* Litchfield 8:l,v:1,2 | ST214 |
| 28S | Human | 28.09.2025 | Almaty city | *S. enterica* Typhimurium 4:i:1,2 | ST19 |
| 29S | Human | 21.09.2025 | Almaty city | *S. enterica* Typhimurium 4:i:1,2 | ST19 |
| 30S | Human | 19.09.2025 | Almaty city | *S. enterica* Muenchen 8:d:1,2 | ST82 |
| 31S | Human | 21.09.2025 | Almaty city | *S. enterica* Muenchen 8:-:1,2 | ST82 |
| 32S | Human | 27.09.2025 | Almaty city | *S. enterica* Typhimurium 4:i:1,2 | ST371<br>8 |
| 36S | Human | 25.10.2025 | Almaty city | *S. enterica* Muenchen 8:d:1,2 | ST82 |
| 37S | Human | 18.11.2025 | Almaty city | *S. enterica* Enteritidis 9:g,m:- | ST11 |
| 40S | Human | 21.11.2025 | Almaty city | *S. enterica* Enteritidis 9:g,m:- | ST11 |
| 41S | Human | 19.12.2025 | Almaty city | *S. enterica* Enteritidis 9:g,m:- | ST11 |

\* Serovar types are reported according to the Kauffmann-White-Le Minor classification scheme [20].
<sup>†</sup> Strain 18S, initially assigned to the genus *Salmonella* based on cultural and biochemical characteristics, was identified as *Proteus mirabilis* by whole-genome sequencing and was subsequently used as a negative control.

### 2.2. DNA extraction

DNA was extracted from pre-decontaminated samples consisting of aqueous bacterial cell suspensions of various concentrations. Bacterial cultures grown overnight at 37°C were inactivated by heating at 100°C for 30 min. Artificially contaminated food samples were treated with sodium merthiolate to a final concentration of 0.01%, followed by incubation at 56 °C for 30 min.

Genomic DNA was extracted using the QIAamp DNA Mini Kit (QIAGEN, Hilden, Germany) according to the manufacturer’s instructions [23]. Purified DNA was eluted in Tris-EDTA (AE) buffer and stored at −20 °C until further use.

### 2.3. PCR and Real-Time PCR conditions

Primer design was performed using the MPrimer 1.4 software package [24]. The specificity of the designed oligonucleotides was evaluated using the BLASTn algorithm against *core_nt* NCBI database [25]. Primers were synthesized by the standard phosphoramidite method using an H6 DNA/RNA synthesizer (K&A Laborgeraete GbR, Germany). Lyophilized primers were dissolved in deionized water to a concentration of 100 μM and stored at −20 °C until use.

Amplification was performed using a Rotor-Gene 6000 thermocycler (QIAGEN, Germany). The reaction mixture had a total volume of 25 μL and contained 10 μL of PCR Mix 1, 10 μL of PCR Mix 2, and 5 μL of template DNA. DNA from the reference strain *S. enteritidis* 3S carrying the target *hilA* gene was used as a positive control. Deionized water without DNA template was used as a negative control.

All real-time PCR reactions were performed in three independent replicates. Results were analysed using Rotor-Gene Q Series Software version 2.3 (QIAGEN, Germany). In accordance with the established assay conditions and the level of background fluorescence, cycle threshold (Ct) values ≤ 37 were considered positive. Samples with Ct values > 37 were considered borderline, whereas samples showing no amplification signal were considered negative.

The developed TaqMan real-time PCR assay employed the following thermal cycling profile: initial denaturation at 95°C for 5 min; 5 cycles of pre-amplification consisting of 95°C for 15 s, 60°C for 20 s, and 72°C for 20 s; followed by 40 amplification cycles with fluorescence acquisition in the FAM/Green channel: 95°C for 15 s, 60°C for 45 s, and 72°C for 20 s.

The nucleotide sequences of the B-*hilA*-F and B-*hilA-R* primers, the TaqMan probe, and the characteristics of the positive control construct are presented in Table 2 (Section B). The screening primers used for multiplex PCR target comparison are provided in Table 2 (Section A).

**Table 2.**
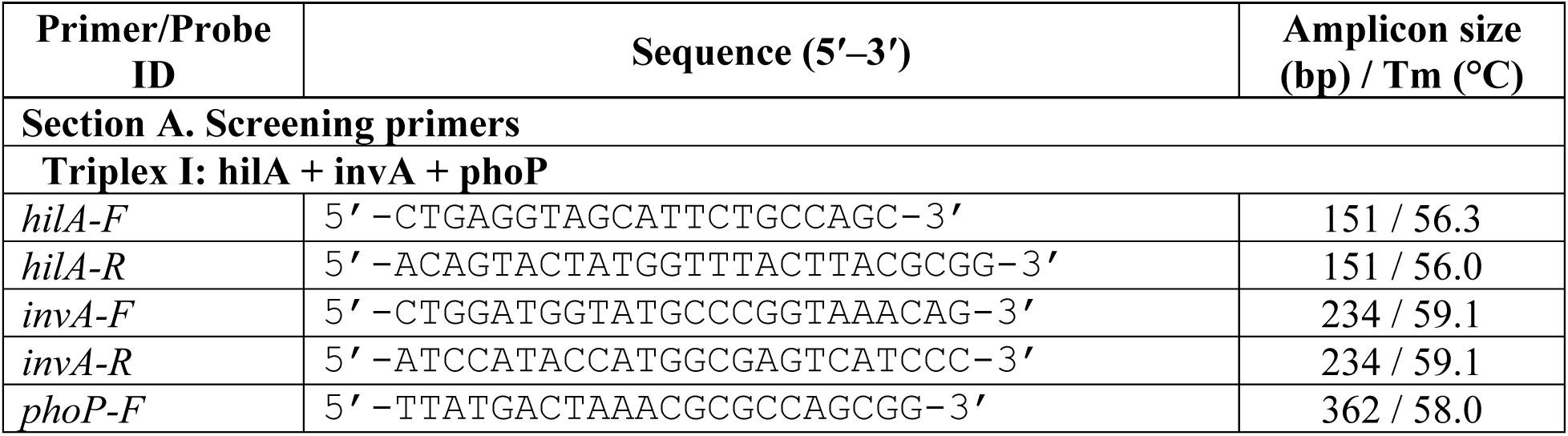
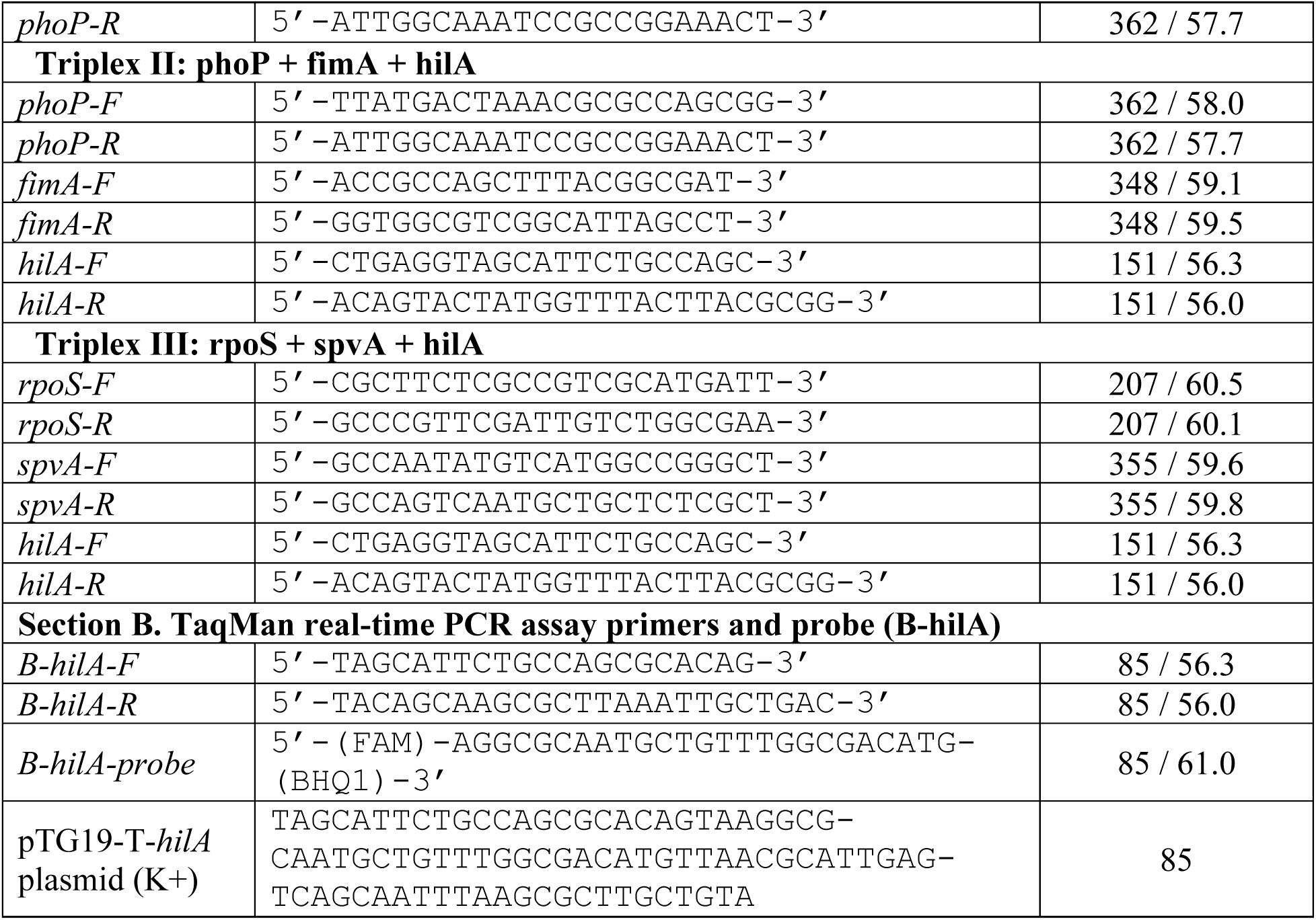
Primers, TaqMan probe, and positive control construct used in this study. Section A: screening primers used in conventional multiplex PCR for target gene comparison. Section B: TaqMan real-time PCR assay oligonucleotides and positive control.

### 2.4. Statistical validation approaches

Quantitative results are presented as the mean ± standard deviation (± SD). Amplification data processing and cycle threshold (Ct) value analysis were performed using Rotor-Gene Q Series Software version 2.3 (QIAGEN, Germany). Additional statistical analyses were conducted using Microsoft Excel 2019 (Microsoft Corp., USA).

To assess assay reproducibility, mean Ct values and standard deviations were calculated across replicates; variation was considered negligible when the coefficient of variation (CV) was below 5%.

To evaluate the analytical performance of the assay, standard curves were generated from the results of ten-fold serial dilutions by plotting Ct values against the logarithm of the concentration of the target matrix DNA or bacterial cells. Amplification efficiency (E) was calculated from the linear regression slope using the formula shown in Eq. 1 [26]:

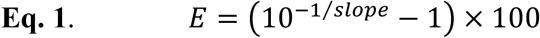

The goodness of fit was assessed using the coefficient of determination (R²). The concentration range over which a linear relationship between Ct values and the logarithm of template concentration was maintained was defined as the assay’s linear dynamic range. In accordance with the MIQE guidelines [26], amplification efficiencies between 90% and 110% and an R² value of at least 0.98 were considered acceptable quality criteria.

### 2.5. Laboratory validation of the assay

Additional validation of the developed TaqMan real-time PCR assay (working name: “NSCEDI Salmonella qPCR”) was performed as a laboratory performance trial at the M. Aikimbayev National Scientific Center for Especially Dangerous Infections (validation protocol dated 11 June 2026). The study included 24 *Salmonella* strains selected from the National Collection of Microorganisms (NCM) of NSCEDI, as well as positive (recombinant plasmid pTG19-T-hilA) and negative (deionized water) control samples.

## 3. Results

### 3.1. Screening of target genes and selection of the optimal primer set

Nucleotide sequences of potential molecular targets for the detection of *Salmonella* spp. were retrieved from the GenBank database (National Center for Biotechnology Information, NCBI, USA). To identify the most universal and specific target, a comparative analysis of the *hilA*, *invA*, *phoP*, *fimA*, *rpoS*, and *spvA* gene sequences was performed. The evaluation considered the degree of gene conservation among different *Salmonella* serovars, including representatives of rare serological groups, as well as their biological relevance and suitability for molecular diagnostics. The *invA* gene was included in the analysis as a classical molecular target that has historically been used for PCR-based detection of *Salmonella* spp. [27].

At the initial stage, three multiplex primer sets were designed for the screening comparison of candidate target genes: *hilA* + *invA* + *phoP*; *phoP* + *fimA* + *hilA*; and *rpoS* + *spvA* + *hilA*. Based on the comparative analysis, the primer set targeting the *hilA* (*hilA*-F/R) and *invA* (*invA*-F/R) genes proved to be the most effective, producing specific amplicons of 151 bp and 234 bp, respectively. The combinations incorporating *phoP*, *fimA*, *rpoS*, and *spvA* targets were excluded from further development because amplification products were absent in one or more isolates representing rare serological groups, indicating insufficient universality across the full isolate panel. The *invA* gene was retained as a secondary marker because it is established as a classical PCR reference target for *Salmonella* detection [27], and its inclusion in the duplex format enabled direct comparison of detection sensitivity with *hilA*. Subsequently, the *hilA*-F/R primers were used as a starting template to design an optimised, shorter inner primer pair (B-*hilA*-F/R) flanking a 85-bp amplicon that accommodated a TaqMan hydrolysis probe, enabling the transition from conventional gel-based PCR to quantitative real-time detection (Table 2, Section B).

Electrophoretic analysis of the amplification products (Figure 1) demonstrated that the *hilA*-F/R primer set consistently detected the target fragment (151 bp) in all 20 isolates confirmed as *Salmonella*, including several representatives of rare serological groups and serovars of diverse origins. In contrast, amplification of the *invA* target fragment (234 bp) was absent in a subset of the tested strains, including several representatives of rare serological groups (lanes 4-20 in the electropherogram shown in Figure 1).

**Figure 1.**
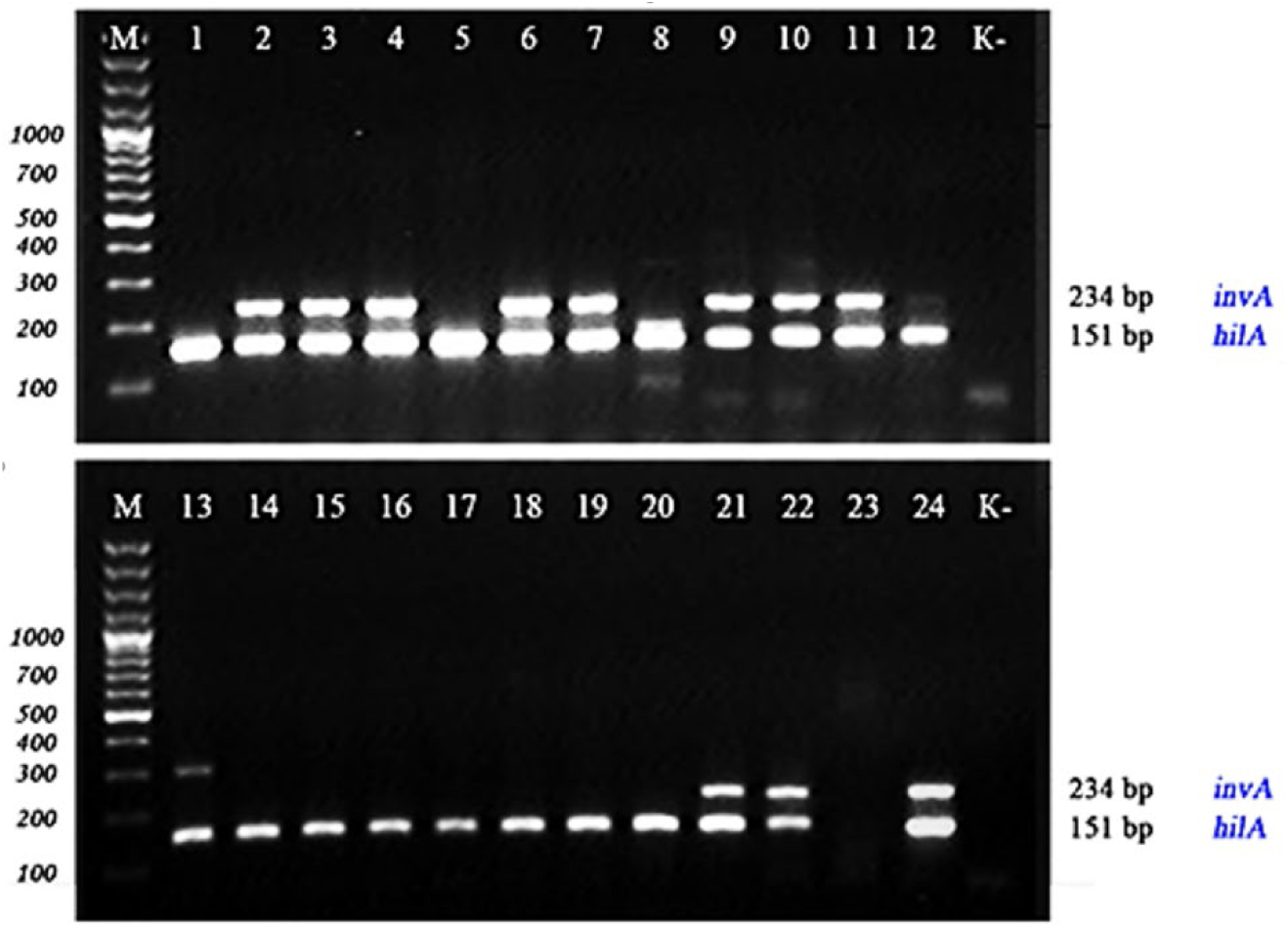
Agarose gel electrophoresis of amplification products obtained with the *inv*A-M1 and *hil*A-M1 primers in a duplex PCR format: M, molecular weight marker; lane 1, *S. typhi 15S*; lane 2, *S. typhimurium 8S*; lanes 3–4, *S. enteritidis 4S, 7S*; lane 5, *S. paratyphi* B 2S; lane 6, *S. dublin 10S*; lane 7, *S. choleraesuis* 22S; lane 8, *S. anatum* 19S; lanes 9–11, *S. enteritidis* 37, 40, 41; lane 12, *S. enteritidis* 34; lane 13, *S. muenchen 50*; lanes 14–20, *Salmonella* other serological groups; lane 21, *S. virchow* 78; lane 22, *S. kentucky* 41; lane 23, *Shigella sonnei* 14; lane 24, positive control; K-, negative control (deionized water).

Based on its superior universality, the *hilA* gene was selected as the primary target for the subsequent development of the TaqMan real-time PCR assay.

### 3.2. Assessment of the analytical specificity of the primer sets

The analytical specificity of the developed *hilA*-M1 and *invA*-M1 primer sets was evaluated by electrophoretic analysis using DNA from *Shigella sonnei 14*, *Shigella flexneri 16*, *Yersinia pseudotuberculosis 2841*, *Y. enterocolitica 5*, *Y. kristensenii 148* and *Vibrio cholerae* ch166, as well as DNA from *S. typhi* 15S, *S. enteritidis* 9S, *S. paratyphi* B 2S, *S. dublin* 10S, *S. infantis 5S*, *S. kottbus 13S* (Figure 2).

**Figure 2.**
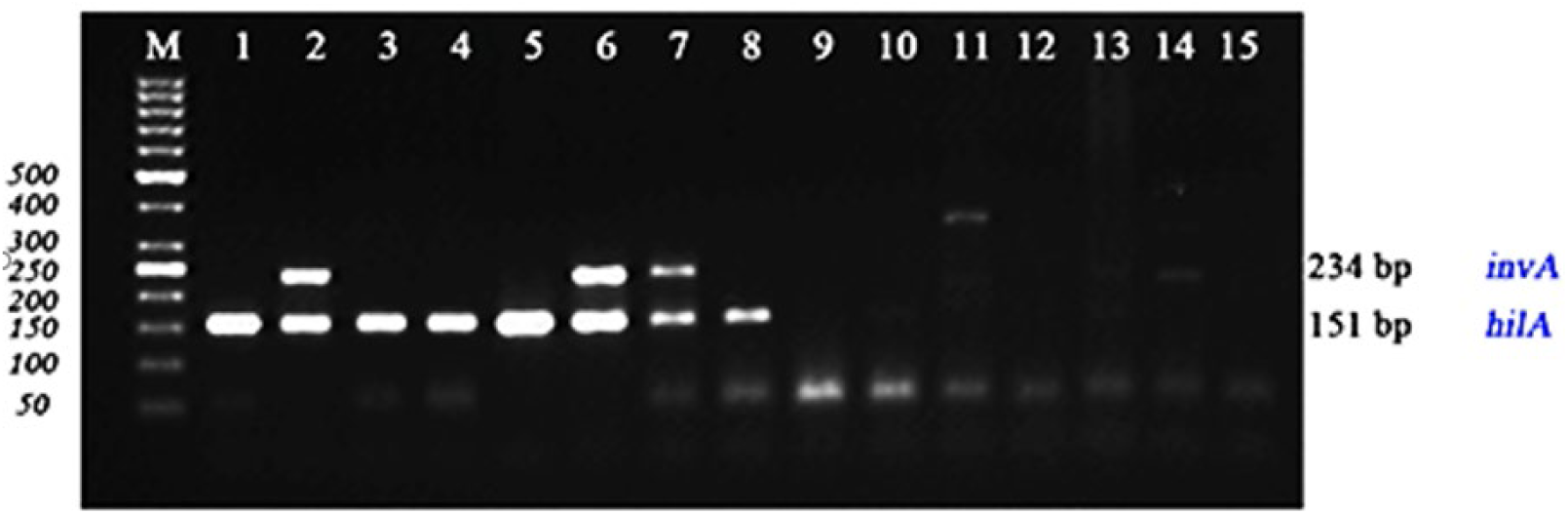
Electrophoretic analysis of PCR products used for the assessment of the analytical specificity of the *hilA*-M1 and *invA*-M1 primer sets. M, DNA molecular weight marker; lane 1, *S. typhi* 15S; lane 2, *S. enteritidis* 9S; lane 3, *S. infantis* 5S; lane 4, *S. kottbus* 13S, lane 5, *S. paratyphi* B 2S; lane 6, *S. dublin* 10S; lane 7, S. *typhimurium* 8S; lane 8, *S. anatum* 19S; lane 9, *Shigella sonnei T13*; lane 10, *Shigella flexneri 16*; lane 11, *Yersinia pseudotuberculosis 2841*; lane 12, *Y. enterocolitica 5*; lane 13, *Y. kristensenii 148*; lane 14, *Vibrio cholerae* ch166; lane 15, negative control (deionized water).

Electrophoretic analysis demonstrated that specific amplification products were detected exclusively in samples containing DNA from members of the genus *Salmonella*. No amplification of the target fragments was observed in any of the non-*Salmonella* species tested. The *hilA*-M1-F/R primers generated a 151-bp amplicon, whereas the *invA*-M1-F/R primers produced a 234-bp amplicon in DNA samples from *Salmonella* positive-control strains. No amplification products were detected in the negative controls, confirming the absence of contamination during PCR setup.

### 3.3. Development and evaluation of the analytical sensitivity of the TaqMan real-time PCR assay for the detection of Salmonella spp

Based on the results of molecular target screening, the *hilA* gene was selected as the primary target for the development of the TaqMan real-time PCR assay. Using the *hilA*-M1-F/R primer pair as a template, a shorter nested primer pair (B-*hilA*-F/R) was designed to generate an 85-bp amplicon containing the binding site for a FAM/BHQ1 TaqMan hydrolysis probe, thereby enabling quantitative real-time detection. The corresponding positive control construct (recombinant plasmid pTG19-T-*hilA*) was constructed to enable run-to-run quality control (Table 2, Section B). The analytical sensitivity of the developed TaqMan real-time PCR assay was evaluated using a series of ten-fold dilutions of genomic DNA from *S. enteritidis* 3S. In all cases, a characteristic relationship between template concentration and cycle threshold (Ct) values was observed: as DNA concentration decreased, Ct values increased proportionally without loss of signal specificity.

Fluorescence amplification curves obtained from the analysis of ten-fold serial dilutions of *S. enteritidis* 3S genomic DNA are shown in Figure 3. The lowest DNA concentration that enabled reproducible detection of the target *hilA* gene fragment in all replicates was 100 fg/μL.

**Figure 3.**
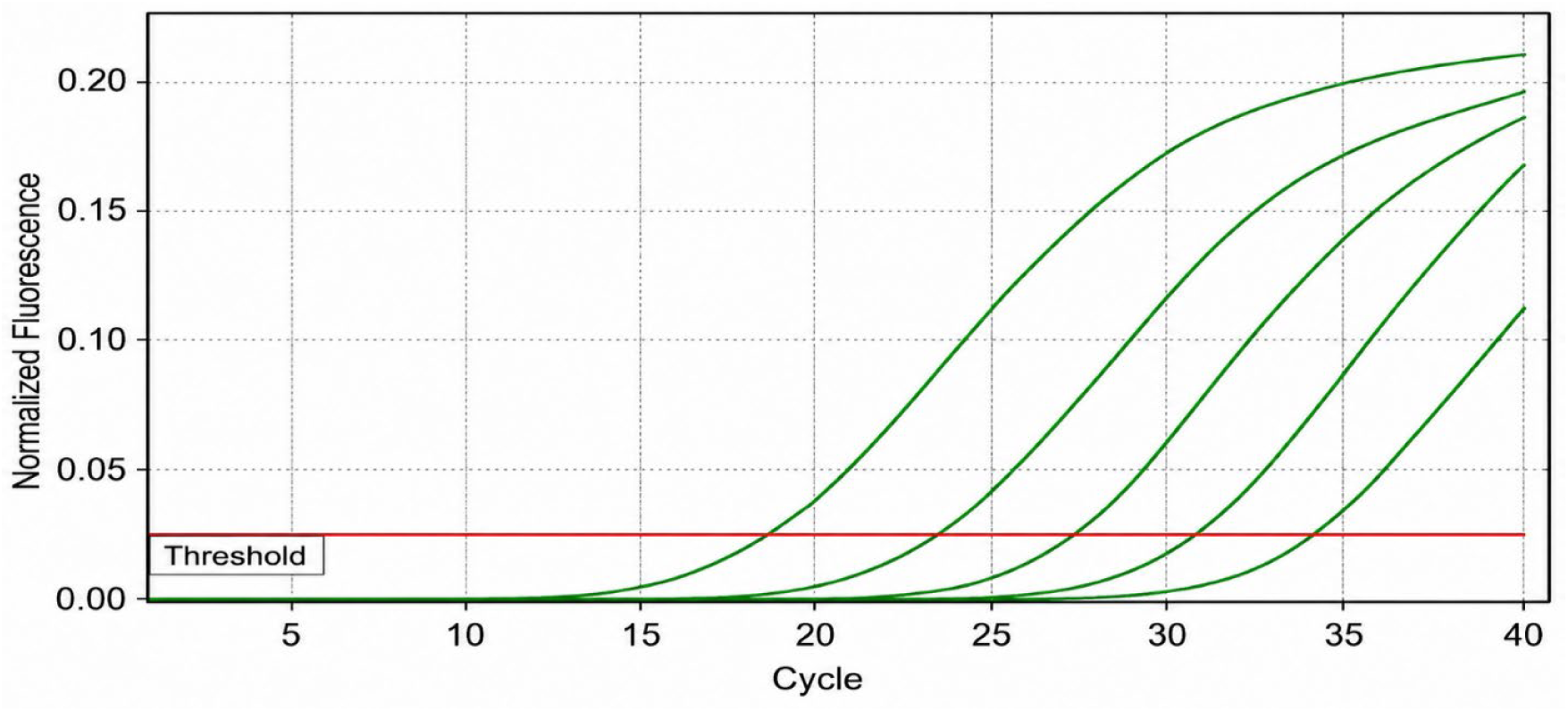
Real-time PCR amplification curves generated from ten-fold serial dilutions of *S. enteritidis* 3S genomic DNA during evaluation of the analytical sensitivity of the developed TaqMan assay.

To conclude, the limit of detection (LOD) of the developed TaqMan real-time PCR assay was 10² bacterial cells/mL when bacterial cell suspensions were used as the test material and 100 fg/μL when genomic DNA was analysed (Figure 3). These results demonstrate the high analytical sensitivity of the assay are comparable to those reported for contemporary real-time PCR methods for the detection of *Salmonella* spp. [28,29].

### 3.4. Evaluation of the analytical specificity of the TaqMan Real-Time PCR Assay

For final validation of the developed assay, an additional assessment of analytical specificity was performed by real-time PCR using an expanded panel of microorganisms, including *Shigella flexneri 16*, *Yersinia pseudotuberculosis 2841*, *Y. pestis NP6*, *Y. enterocolitica 5*, *Y. kristensenii 148*, *Bacillus anthracis 23*, *Vibrio cholerae ch166* and *Francisella tularensis FT24*. Tree of these organisms: *Y. pestis NP6*, *B. anthracis 23* and *F. tularensis FT24*, were not included in the agarose gel electrophoresis screening described above because of biosafety requirements associated with handling these microorganisms.

No specific amplification was detected in any of the non-target microorganisms tested. A positive fluorescence signal was observed exclusively in samples containing DNA from *S. enterica* Enteritidis, which served as the positive control. Ct values were undetermined for all eight non-target microorganisms, indicating the absence of cross-amplification. Together with the negative results obtained during the gel electrophoresis screening (Figure 2), these findings demonstrate complete analytical specificity of the developed assay for bacteria of the genus *Salmonella*.

### 3.5. Evaluation of assay performance using artificially contaminated food samples

The performance of the developed TaqMan real-time PCR assay was evaluated using artificially contaminated food samples. Milk and carrot were used as model food matrices and were spiked with known concentrations of *S. enteritidis* 3S cell suspensions. Following DNA extraction, the samples were analysed by real-time PCR according to the developed protocol.

Specific amplification of the target *hilA* gene fragment was detected in all artificially contaminated samples. Cycle threshold (Ct) values ranged from 18.4 to 22.7, indicating efficient detection of *Salmonella* DNA in the presence of food-matrix components. No amplification was observed in the negative-control samples. These results indicate the absence of substantial inhibitory effects of the tested food matrices on PCR amplification and confirm the suitability of the developed assay for the detection of *Salmonella* spp. in food products.

### 3.6. Evaluation of assay performance using bacterial isolates

The diagnostic performance of the developed TaqMan real-time PCR assay was evaluated using a collection of bacterial strains obtained from the Laboratory of the NCM at the NSCEDI. The collection comprised 25 strains, including 24 *Salmonella* spp. strains and one *Proteus mirabilis* strain. Following DNA extraction, all samples were analysed using the B-hilA-F/R primers and the specific TaqMan probe.

Specific amplification of the target *hilA* gene fragment was detected in all 24 strains confirmed as *Salmonella*. Cycle threshold (Ct) values ranged from 14.04 to 38.68 (Table 3). For most positive strains, Ct values did not exceed 24; however, one strain (25S, *S. enterica* Dublin) yielded a signal above the positivity threshold (Ct = 38.68), which was interpreted as a weakly positive result. No amplification was detected for strain 18S, which had been identified by whole-genome sequencing as *P. mirabilis*. The positive control sample (recombinant plasmid pTG19-T-hilA) showed consistent amplification (Ct = 11.91), whereas no amplification products were detected in the negative control containing nuclease-free water. Amplification curves obtained from the analysis of the bacterial isolates are shown in Figure 4.

**Figure 4.**
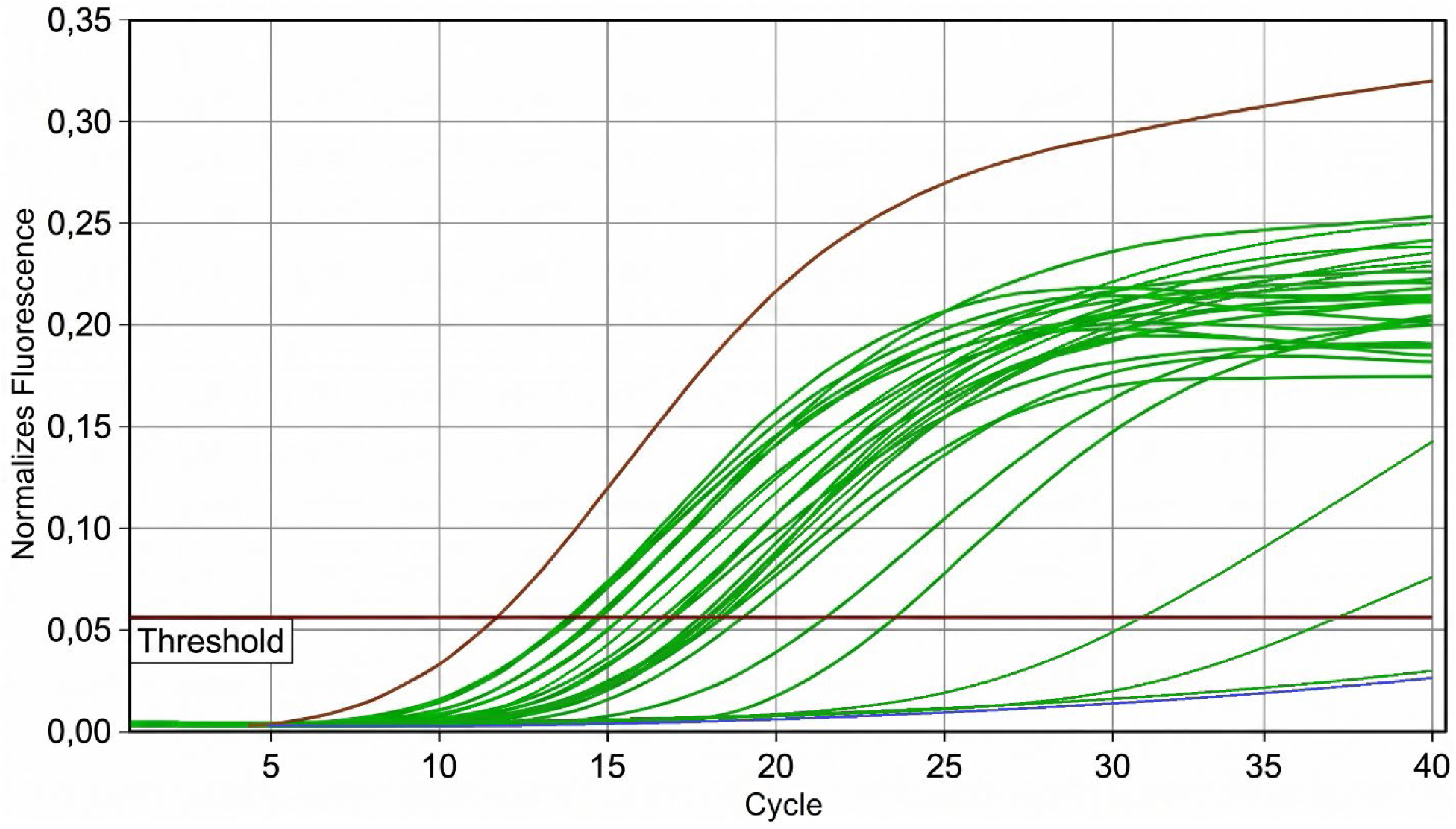
Amplification curves obtained from the analysis of 24 *Salmonella* spp. isolates and one *Proteus mirabilis* isolate using the TaqMan real-time PCR assay targeting the *hilA* gene. The red curve represents the positive control sample (pTG19-T-*hilA* plasmid), and the horizontal line indicates the fluorescence threshold.

**Table 3.** Results of TaqMan real-time PCR detection of *Salmonella* spp. isolates targeting the *hilA* gene.

| Strain | Species and serovar | Ct |
| --- | --- | --- |
| 2S | *S. enterica* Paratyphi B | 14,04 |
| 5S | *S. enterica* Infantis | 15,60 |
| 6S | *S. enterica* Enteritidis | 14,87 |
| 8S | *S. enterica* Typhimurium | 15,07 |
| 10S | *S. enterica* Dublin | 18,69 |
| 13S | *S. enterica* Kottbus | 20,32 |
| 15S | *S. enterica* Typhi | 19,92 |
| 16S | *S. enterica* Anatum | 18,12 |
| 17S | *S. enterica* Enteritidis | 17,93 |
| 18S | *P. mirabilis* | – |
| 20S | *S. enterica* Typhimurium | 15,92 |
| 23S | *S. enterica* Choleraesuis | 18,23 |
| 24S | *S. enterica* Choleraesuis | 19,47 |
| 25S | *S. enterica* Dublin | 38,68 |
| 26S | *S. enterica* Typhimurium | 17,01 |
| 27S | *S. enterica* Litchfield | 15,58 |
| 28S | *S. enterica* Typhimurium | 17,10 |
| 29S | *S. enterica* Typhimurium | 31,28 |
| 30S | *S. enterica* Muenchen | 14,16 |
| 31S | *S. enterica* Muenchen | 18,63 |
| 32S | *S. enterica* Typhimurium | 23,72 |
| 36S | *S. enterica* Muenchen | 14,07 |
| 37S | *S. enterica* Enteritidis | 14,86 |
| 40S | *S. enterica* Enteritidis | 14,25 |
| 41S | *S. enterica* Enteritidis | 14,94 |

## 4. Discussion

One of the principal findings of this study was the demonstration of the broader distribution and higher diagnostic utility of the *hilA* gene as a diagnostic marker compared with the widely used *invA* gene for PCR-based detection of *Salmonella*. Unlike *invA*, which was absent in a subset of the isolates examined, including representatives of rare serological groups, the *hilA* gene was detected in all tested *Salmonella* strains. These findings are consistent with previously published studies that have identified *hilA* as a highly conserved regulator of the SPI-1 pathogenicity island and a promising target for the universal detection of *Salmonella* [30–33].

Genomic characterization of a subset of the strain collection, performed by whole-genome sequencing as part of a parallel study [22], provides additional insight into the differences in the universality of the molecular targets evaluated. The *hilA* gene is located within the SPI-1 pathogenicity island, a conserved chromosomal element of *S. enterica* that is present irrespective of the plasmid content of a given strain. In contrast, the *spvA* gene, one of the six candidate targets initially considered in this study, is associated with the virulence genes *spvB* and *spvC*, which are located on a large virulence plasmid. This plasmid is not present in all isolates and may be lost or replaced during strain adaptation by an alternative plasmid carrying a similar repertoire of genes [22, 34]. A similar dependence on plasmid content may also apply to other candidate targets that did not demonstrate complete universality during the screening stage.

Thus, the chromosomal localization of *hilA* and its regulatory role in controlling the expression of virulence genes within the SPI-1 pathogenicity island [30–33] provide a plausible molecular-genetic explanation for its greater conservation relative to plasmid-associated markers and further support its selection as a target for the development of a universal diagnostic assay.

Of particular importance is the fact that the broad applicability of the proposed primer set was confirmed in this study across both widely distributed serovars of the pathogen (*S. enterica* Enteritidis, *S. enterica* Choleraesuis, and *S. enterica* Dublin) and representatives of less common serovars (*S. enterica* Newport, *S. enterica* Infantis, *S. enterica* Kottbus, *S. enterica* Anatum, and *S. enterica* Typhi).

According to international studies, the diversity of circulating non-typhoidal *Salmonella* serovars continues to increase. For example, an analysis of data from 15 countries in the Americas identified representatives of 131 *Salmonella* serovars in poultry populations [35]. Likewise, in countries of the European Union, in addition to *S. enterica* Enteritidis and Typhimurium, the monophasic variant of *S. enterica* Typhimurium, *S. enterica* Infantis, and *S. enterica* Derby are widely distributed and are also among the most frequently detected serovars in African countries [36]. As the serological diversity of circulating *Salmonella* strains continues to increase, the use of a universal molecular target is becoming increasingly important for ensuring reliable laboratory diagnosis and minimizing the risk of false-negative results associated with atypical or less common serovars.

From a practical perspective, the use of a single highly conserved molecular target enables simplification of assay design while maintaining high analytical sensitivity and specificity. The developed TaqMan real-time PCR assay exhibited a limit of detection of 10² bacterial cells/mL when bacterial cell suspensions were used as the test material and 100 fg/μL when genomic DNA was analysed. This level of sensitivity is comparable to that reported for contemporary real-time PCR assays used for the laboratory detection of *Salmonella* spp. [28,29,33,37,38].

Another important finding of this study is the high analytical specificity of the developed assay. No cross-amplification was observed when DNA samples from *Shigella flexneri 16*, *Shigella sonnei 14*, *Yersinia pestis NP6*, *Y. pseudotuberculosis 2841*, *Y. enterocolitica 5*, *Y. kristensenii 148*, *Bacillus anthracis 23*, *Vibrio cholerae ch166* and *Francisella tularensis FT24* were tested. Additional independent evidence supporting the specificity of the assay was provided by strain 18S, which had initially been assigned to the genus *Salmonella* based on cultural and biochemical characteristics but was subsequently identified as *P. mirabilis* by whole-genome sequencing. The developed assay did not produce specific amplification with DNA from this strain, consistent with the whole-genome sequencing results and demonstrating the ability of the method to distinguish *Salmonella* from other members of the family Enterobacteriaceae. Taken together, these findings support the appropriateness of selecting the *hilA* gene as a diagnostic target.

The practical applicability of the assay was further demonstrated using artificially contaminated food samples, in which specific amplification was detected in all positive samples containing cultured *Salmonella* bacteria. Overall, the results indicate that the developed assay possesses high analytical sensitivity and specificity and has considerable potential for application in the laboratory detection of *Salmonella*, food safety monitoring, and epidemiological surveillance, particularly in the context of the increasing diversity of circulating *Salmonella* serovars, including representatives of rare serological groups.

### Limitations of this study

The present study has several limitations that should be considered when interpreting the results.

First, the performance characteristics of the developed assay were evaluated using a limited collection of *Salmonella* isolates obtained predominantly from Kazakhstan. Although the study included representatives of multiple serovars, including rare serological groups, this collection does not encompass the full genetic diversity of the genus *Salmonella*. Further studies involving a larger number of samples from diverse sources, including food, veterinary, environmental, and clinical specimens, are required for comprehensive validation and broader assessment of assay performance.

Second, the analytical sensitivity and specificity of the assay were assessed using a limited panel of heterologous microorganisms. Further studies involving a larger number of samples from diverse sources, including food, veterinary, and environmental specimens, are required for comprehensive validation and broader assessment of assay performance.

Third, the study was designed for the qualitative detection of *Salmonella* spp. and did not include quantitative assessment of bacterial load or PCR-based determination of the serovar identity of detected isolates.

## 5. Conclusions

In the present study, a TaqMan real-time PCR assay for the detection of *Salmonella* spp. based on the *hilA* gene was developed and validated. Comparative evaluation of six candidate molecular targets (*hilA*, *invA*, *phoP*, *fimA*, *rpoS*, and *spvA*) demonstrated that the *hilA* gene exhibited the broadest coverage and enabled the detection of all *Salmonella* strains included in the study, including representatives of rare serological groups.

The developed assay demonstrated high analytical sensitivity, with limits of detection of 10² bacterial cells/mL and 100 fg/μL of genomic DNA, as well as high analytical specificity, with no cross-reactivity observed when tested against a panel of heterologous microorganisms. The performance of the assay was further confirmed using artificially contaminated food samples.

These findings indicate that the *hilA* gene represents a promising universal molecular target for the detection of *Salmonella* spp. Furthermore, the developed TaqMan real-time PCR assay has potential applications in the laboratory detection of *Salmonella*, surveillance of pathogen circulation, and strengthening of epidemiological monitoring programs, including within the framework of the One Health approach for integrated monitoring of *Salmonella* in human populations, animals, food products, and environmental reservoirs.

## Institutional Review Board Statement

The study was conducted in accordance with the principles of the Declaration of Helsinki and was approved by the Local Bioethics Committee of the M. Aikimbayev National Scientific Center for Especially Dangerous Infections (Protocol No. 2, dated 12 February 2024; Section 2.1).

## References

1. Cherenova LP, Zakayev KY, Matsui AV, Cherenova VK. Clinical characteristics of intestinal mixed infections. Perm Medical Journal. 2023 Jun 2;40(2):29–38.DOI: 10.17816/pmj40229-38.

2. Gazezova, S.; Nabirova, D.; Waltenburg, M.; Rakhimzhanova, M.; Smagul, M.; Kasabekova, L.; Horth, R. Salmonellosis outbreak associated with the consumption of food at a wedding in an urban restaurant in Kazakhstan: a retrospective cohort study. BMC Infect. Dis. 2024, 24, 1474. DOI: 10.1186/s12879-024-10382-4.

3. Woh, P.Y.; Yeung, M.P.S.; Goggins, W.B.; Lo, N.; Wong, K.T.; Chow, V.; Chau, K.Y.; Fung, K.; Chen, Z.; Ip, M. Genomic Epidemiology of Multidrug-Resistant Nontyphoidal. *Salmonella* in Young Children Hospitalized for Gastroenteritis. Microbiol. Spectr. 2021, 9, e00248–21. DOI: 10.1128/spectrum.00248-21.

4. Majowicz, S.E.; Musto, J.; Scallan, E.; Angulo, F.J.; Kirk, M.; O’Brien, S.J.; Jones, T.F.; Fazil, A.; Hoekstra, R.M.; International Collaboration on Enteric Disease ‘Burden of Illness’ Studies. The global burden of nontyphoidal *Salmonella* gastroenteritis. Clin. Infect. Dis. 2010, 50, 882–889. DOI: 10.1086/650733.

5. Stanaway, J.D.; Parisi, A.; Sarkar, K.; Blacker, B.F.; Reiner, R.C.; Hay, S.I.;, et al. The global burden of non-typhoidal *Salmonella* invasive disease: a systematic analysis for the Global Burden of Disease Study 2017. Lancet Infect. Dis. 2019, 19, 1312–1324. DOI: 10.1016/S1473-3099(19)30418-9.

6. Ranjan, A.; Chandna, M.; Stevens, N.J.; Kandil, J.; Dinh, B.; Kuhn, M.; Mian, N.; Tran, B.; Hamid, A.; Kim, P.; Desin, T.S. *Salmonella* infections: global trends and emerging challenges. Microorganisms 2026, 14, 816. DOI: 10.3390/microorganisms14040816.

7. Chae S.J., Yun Y.S., Yoo C.K., Chung G.T., Lee D.Y. First Report of Salmonella Serotype Tilene Infection in Korea. Ann Clin Microbiol. 2016;19(1):24–27. doi:10.5145/ACM.2016.19.1.24.

8. European Food Safety Authority; European Centre for Disease Prevention and Control. The European Union summary report on antimicrobial resistance in zoonotic and indicator bacteria from humans, animals and food in 2021–2022. EFSA J. 2024, 22, e8583. 10.2903/j.efsa.2025.9237

9. Nogaybek MD, Baigenzheeva RK. Analysis of Salmonellosis Incidence in Kazakhstan During 2018–2022. Nauka i Mirovozzrenie (Science and Worldview). 2024;1:13–15. (available from https://cyberleninka.ru/article/n/analiz-zabolevaemosti-salmonellezom-v-kazahstane-za-2018-2022-gody/viewer)

10. Baimuratova, M.; Abdul, B.A.; Ryskulova, A.; Tugulbayeva, A.; Jumatova, U.; Abdusallamova, Z. Microbiological monitoring in the system of epidemiological surveillance of salmonellosis in the children’s population of Almaty city. Наука и здравоохранение 2021, 23, 121–130. DOI: 10.34689/SH.2021.23.3.014.

11. Grjibovski, A.M.; Kosbayeva, A.; Menne, B. The effect of ambient air temperature and precipitation on monthly counts of salmonellosis in four regions of Kazakhstan, Central Asia, in 2000–2010. Epidemiol. Infect. 2014, 142, 608–615. DOI: 10.1017/S095026881300157X.

12. Karaaslan, A.; Soysal, A.; Kadayifci, E.K.; Yakut, N.; Akkoc, G.; Demir, S.O.; Atici, S.; Söyletir, G.; Bakir, M. Salmonella gastroenteritis in children. J. Infect. Dev. Ctries 2022, 16, 1757–1761. DOI: 10.3855/jidc.17042.

13. Sapega E.Yu., Butakova L.V., Trotsenko O.E. Modern epidemiological features of viral acute intestinal infections in children and adolescents of Sakhalin region. Journal Infectology. 2024;16(3):123–132. doi:10.22625/2072-6732-2024-16-3-123-132.

14. Andrews, J.R.; Ryan, E.T. Diagnostics for invasive *Salmonella* infections: current challenges and future directions. Vaccine 2015, 33 (Suppl. 3), C8–C15. DOI: 10.1016/j.vaccine.2015.02.030.

15. Espy, M.J.; Uhl, J.R.; Sloan, L.M.; Buckwalter, S.P.; Jones, M.F.; Vetter, E.A.;, et al. Real-time PCR in clinical microbiology: applications for routine laboratory testing. Clin. Microbiol. Rev. 2006, 19, 165–256. DOI: 10.1128/CMR.19.1.165-256.2006.

16. Galán, J.E.; Ginocchio, C.; Costeas, P. Molecular and functional characterization of the *Salmonella* invasion gene *invA*: homology of InvA to members of a new protein family. J. Bacteriol. 1992, 174, 4338–4349. DOI: 10.1128/jb.174.13.4338-4349.1992.

17. Ohl, M.E.; Miller, S.I. *Salmonella*: a model for bacterial pathogenesis. Annu. Rev. Med. 2001, 52, 259–274. DOI: 10.1146/annurev.med.52.1.259.Factors of *Salmonella* 2001, 52, 259–274. DOI: 10.1146/annurev.med.52.1.259.

18. Boddicker, J.D.; Knosp, B.M.; Jones, B.D. Transcription of the *Salmonella* invasion gene activator, *hilA*, requires HilD activation in the absence of negative regulators. J. Bacteriol. 2003, 185, 525–533. DOI: 10.1128/JB.185.2.525-533.2003.

19. Adesiji, Y.O.; Deekshit, V.K.; Odunola, R.A.; Karunasagar, I.; Daodu, O.B.; Ahmad, A.M. Virulence-encoding genes conserved in *Salmonella* isolated from humans, poultry, and seafood. J. Trop. Med. 2025, 2025, 1139253. DOI: 10.1155/jotm/1139253.

20. Andreev IА, Baranov DA, Vecherkin VA, Ptitsyn VA, Koryashkin PV, Gagloev VМ. Salmonellosis osteomyelitis of the pelvic bones in adolescent: a case report. Russian Journal of Pediatric Surgery, Anesthesia and Intensive Care. 2024 Jul 16;14(2):267–76. DOI: 10.17816/psaic1782.

21. Grimont, P.A.D.; Weill, F.-X. Antigenic Formulae of the Salmonella Serovars, 9th ed.; WHO Collaborating Centre for Reference and Research on Salmonella, Institut Pasteur: Paris, France, 2007.

22. Yessimseit DT, Rysbekova AK, Zhumadilova ZB, Abdeliyev BZ, Kassenova AK, Tukhanova NB, Abdrakhmanova AK, Mereke A, Agzam SD, Nurpeisova AS, Nissanova R. Genetic and epigenetic diversity of *Salmonella enterica* isolates from Kazakhstan from clinical and veterinary sources. bioRxiv. 2026:2026–06. DOI: 10.64898/2026.06.01.729238.

23. QIAGEN. QIAamp DNA Mini and Blood Mini Handbook; QIAGEN: Hilden, Germany, 2020. (https://dna.uga.edu/wp-content/uploads/sites/51/2013/12/QIA-amp-DNA-Blood-Kits-Handbook.pdf)

24. Shen, Z.; Qu, W.; Wang, W.; Lu, Y.; Wu, Y.; Li, Z.; Hang, X.; Wang, X.; Zhao, D.; Zhang, C. MPrimer: a program for reliable multiplex PCR primer design. BMC Bioinform. 2010, 11, 143. DOI: 10.1186/1471-2105-11-143.

25. Altschul, S.F.; Gish, W.; Miller, W.; Myers, E.W.; Lipman, D.J. Basic local alignment search tool. J. Mol. Biol. 1990, 215, 403–410. DOI: 10.1016/S0022-2836(05)80360-2.

26. Bustin, S.A.; Benes, V.; Garson, J.A.; Hellemans, J.; Huggett, J.; Kubista, M.; Mueller, R.; Nolan, T.; Pfaffl, M.W.; Shipley, G.L.;, et al. The MIQE guidelines: minimum information for publication of quantitative real-time PCR experiments. Clin. Chem. 2009, 55, 611–622. DOI: 10.1373/clinchem.2008.112797.

27. Rahn, K.; De Grandis, S.A.; Clarke, R.C.; McEwen, S.A.; Galán, J.E.; Ginocchio, C.; Curtiss, R.; Gyles, C.L. Amplification of an *invA* gene sequence of *Salmonella typhimurium* by polymerase chain reaction as a specific method of detection of *Salmonella*. Mol. Cell. Probes 1992, 6, 271–279. DOI: 10.1016/0890-8508(92)90002-F.

28. Malorny, B.; Paccassoni, E.; Fach, P.; Bunge, C.; Martin, A.; Helmuth, R. Diagnostic real-time PCR for detection of *Salmonella* in food. Appl. Environ. Microbiol. 2004, 70, 7046–7052. DOI: 10.1128/AEM.70.12.7046-7052.2004.

29. Postollec, F.; Falentin, H.; Pavan, S.; Combrisson, J.; Sohier, D. Recent advances in quantitative PCR (qPCR) applications in food microbiology. Food Microbiol. 2011, 28, 848–861. DOI: 10.1016/j.fm.2011.02.008.

30. Bajaj, V.; Hwang, C.; Lee, C.A. *hilA* is a novel *ompR/toxR* family member that activates the expression of *Salmonella typhimurium* invasion genes. Mol. Microbiol. 1995, 18, 715–727. DOI: 10.1111/j.1365-2958.1995.mmi_18040715.x.

31. Pathmanathan, S.G.; Cardona-Castro, N.; Sánchez-Jiménez, M.M.; Correa-Ochoa, M.M.; Puthucheary, S.D.; Thong, K.L. Simple and rapid detection of *Salmonella* strains by direct PCR amplification of the *hilA* gene. J. Med. Microbiol. 2003, 52, 773–776. DOI: 10.1099/jmm.0.05118-0.

32. Guo, X.; Chen, J.; Beuchat, L.R.; Brackett, R.E. PCR detection of *Salmonella enterica* serotype Montevideo in and on raw tomatoes using primers derived from *hilA*. Appl. Environ. Microbiol. 2000, 66, 5248–5252. DOI: 10.1128/AEM.66.12.5248-5252.2000.

33. Cardona-Castro, N.; Sánchez-Jiménez, M.; Lavalett, L.; Múñera-Jaramillo, M.; Correa-Ochoa, M.M. Validation of a PCR for diagnosis of typhoid fever and salmonellosis by amplification of the *hilA* gene in clinical samples from Colombian patients. J. Med. Microbiol. 2009, 58, 913–919. DOI: 10.1099/jmm.0.008797-0.

34. Tasmin, R.; Gulig, P.A.; Parveen, S. Detection of virulence plasmid-encoded genes in *Salmonella* Typhimurium and *Salmonella* Kentucky isolates recovered from commercially processed chicken carcasses. J. Food Prot. 2019, 82, 1364–1368. DOI: 10.4315/0362-028X.JFP-18-552.

35. Diaz, D.; Hernandez-Carreño, P.E.; Velazquez, D.Z.;, et al. Prevalence, main serovars, and antimicrobial resistance profiles of non-typhoidal *Salmonella* in poultry samples from the Americas: a systematic review and meta-analysis. Authorea (preprint) 2021. DOI: 10.22541/au.163253387.70584910/v1.

36. European Food Safety Authority; European Centre for Disease Prevention and Control. The European Union One Health 2021 zoonoses report. EFSA J. 2022, 20, e7666. DOI: 10.2903/j.efsa.2022.7666.

37. Elizaquível, P.; Aznar, R.; Sánchez, G. Recent developments in the use of viability dyes and quantitative PCR in the food microbiology field. J. Appl. Microbiol. 2014, 116, 1–13. DOI: 10.1111/jam.12365.

38. Wilson, I.G. Inhibition and facilitation of nucleic acid amplification. Appl. Environ. Microbiol. 1997, 63, 3741–3751. DOI: 10.1128/aem.63.10.3741-3751.1997.

